# Mapping the Neural Control of Force and Impedance of Wrist Movements Using Robotics and fMRI

**DOI:** 10.1101/2025.01.03.631219

**Authors:** Kristin Schmidt, Fabrizio Sergi

## Abstract

While robots are becoming increasingly valuable tools in neurorehabilitation, our limited understanding of the brain’s response during human-robot interaction tasks restricts advancements in training programs to restore neural pathways after injury. Co-contraction is characteristic of several neuro-muscular disorders, such as stroke and cerebral palsy, and it is often targeted by assessments or rehabilitation programs. Despite its importance, the neural mechanisms underlying co-contraction remain poorly understood. To address this gap, this study investigates the neural substrates of muscle co-contraction via functional magnetic resonance imaging (fMRI) during a dynamic motor task with an MR-compatible wrist robot.

To establish suitable fMRI experimental conditions, we first conducted a behavioral study assessing muscle activity during a wrist-pointing task with four participants. Participants reached toward a target while experiencing four main perturbation conditions (no force, divergent force, constant force up, and constant force down), designed to elicit distinct force and impedance responses. Following this behavioral validation, five additional participants performed the wrist-pointing task during fMRI. Our results suggest localization of force and impedance control within the cortico-thalamic-cerebellar network. These findings provide new insights into the neural mechanisms of co-contraction, supporting the development of neurorehabilitation paradigms.

## I. Introduction

Motor control is essential for performing skilled movements in daily life, yet disruptions in this ability are associated with several neuromuscular conditions, such as stroke and cerebral palsy [1]. These neuromuscular disorders often feature altered muscle activation strategies, such as excessive co-contraction of agonist and antagonist muscles. This lack of precise regulation of muscle groups impairs movement efficiency and increases fatigue, making everyday activities more challenging. As a result, physical therapists sometimes target a reduction in muscle co-contraction as a goal for rehabilitation [1]. However, current therapeutic approaches to regulating muscle co-contraction are limited by our lack of understanding of how muscle co-contraction is controlled at the neural level. Addressing this gap is critical for identifying therapeutic targets and developing effective interventions to optimize muscle coordination and restore motor function.

Co-contraction plays an important role in maintaining joint stability [2], resisting external disturbances during movement [3], and may also facilitate the learning of new skills [4]. However, it is a metabolically demanding control strategy and is generally reserved for tasks that demand increased joint stiffness, such as those that are unpredictable or unstable. Conversely, tasks that involve learnable dynamics allow the central nervous system to predict dynamics with an accurate internal model, enabling a more efficient control strategy that relies on the reciprocal activation of agonist muscles. While co-contraction and reciprocal activation are distinct motor strategies, they often occur simultaneously, as we have shown in our previous work in the wrist joint [5], [6], necessitating careful selection of task conditions to isolate their control.

In this work, we first validate a set of experimental task conditions in a wrist-pointing task designed to decouple co-contraction and reciprocal activation, as estimated by electromyography (EMG). This EMG validation is essential for ensuring that the task conditions reliably differentiate between these mechanisms, a crucial step for distinguishing their underlying neural processes. Following this behavioral validation, we conduct an fMRI study using the same task conditions to investigate the neural correlates associated with torque and stiffness control. This work provides preliminary insights into the neural mechanisms responsible for muscle co-contraction and reciprocal activation during dynamic motor tasks, with potential implications for improving rehabilitation strategies.

## II. Materials and Methods

In this work, we present findings from two studies, a behavioral study and a neuroimaging study, with similar methodologies; the primary distinction between the two studies is that the behavioral study was conducted in a mock MRI scanner with EMG, while the neuroimaging study took place in a real MRI scanner without EMG. In both studies, participants performed wrist-pointing movements using the MR-SoftWrist, an MR-compatible wrist robot, while lying supine (Fig. 1).

**Fig 1.**
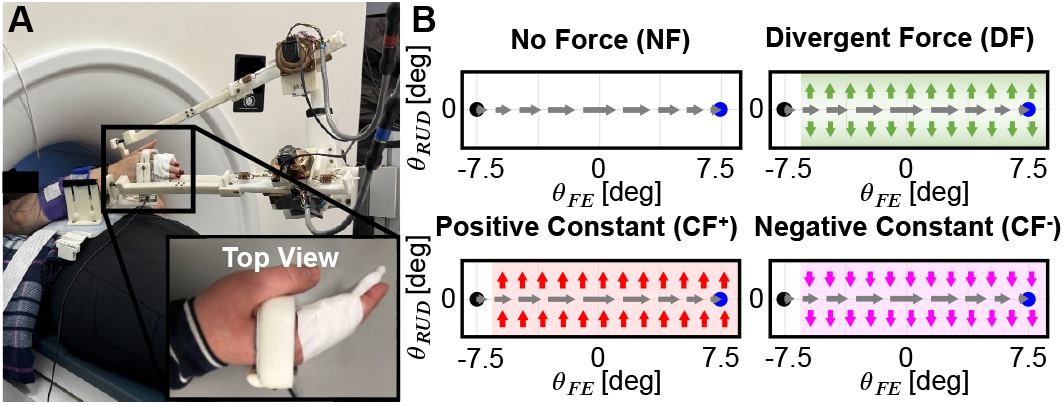
A. MR-SoftWrist pictured in its operating condition. The enlarged image shows a top view of the handle with the paddle insert and tape around the participant’s fingers. B. Experimental force conditions.

### A. Participants

Four healthy young adults (2 male, 27.5 *±* 3.11 years) were recruited for the behavioral study, and five healthy young adults (3 male, 27.2 *±* 3.19 years) were recruited for the neuroimaging study. Participants had no history of neuromuscular disorders. This study was approved by the Institutional Review Board of the University of Delaware, IRB no. 1936400-3.

### B. Experimental Setup

Participants moved the handle of the MR-SoftWrist with their right wrist in flexion-extension (FE) and radial-ulnar deviation (RUD). Their hand was secured in the robot’s handle, which enclosed the knuckles without limiting finger movement. A small paddle was inserted into the handle to promote a slightly flexed finger posture and reduce extraneous finger muscle contraction, and the participant’s fingers were taped to the paddle (Fig. 1A). Participants were instructed to focus on making wrist movements while keeping fingers relaxed to minimize additional confounds due to finger muscle contraction. Their forearm was supported by a rigid arm support placed on their stomach, with additional padding for shoulder and upper arm support. Participants viewed the task on a monitor through a pair of glasses with angled mirrors in the behavioral study or through an angled mirror attached to the imaging head coil in the neuroimaging study.

In all tasks, the goal was to move in a straight line between two circular targets. For horizontal tasks, these were located at (*±* 7.5,0) deg, and for vertical tasks, these were located at (0, *±* 7.5) deg, relative to the neutral hand posture in the FE and RUD axes, respectively. Each trial began with the target changing color from black to blue. After the participant moved the cursor 1 deg away from the starting target, the cursor was hidden to reduce corrections on real-time feedback. At the end of each trial, speed feedback was displayed by a change in target color to red if too slow (*>*0.45 s), green if too fast (*<*0.2 s), or black if within the desired time window. Error feedback was also displayed via a yellow cursor at the position of the maximum deviation from the straight line trajectory.

During these movements, the robot applied forces perpendicular to the primary movement direction. The main force conditions used in this study were no force (NF), divergent force (DF), constant force up (CF^+^), and constant force down (CF^*−*^) (Fig. 1B). NF was intended to assess baseline performance. In DF, the robot applied RUD perturbation torques proportional to the RUD wrist angle, up to a maximum torque, *τ*_*m*_, such that *τ*_*RUD*_ = min(*k*_*v*_*θ*_*RUD*_, *τ*_*m*_), with *k*_*v*_ = 0.085 Nm/deg and *τ*_*m*_ = 1 Nm. This unstable condition was designed to up-regulate muscle co-contraction [3]. In the CF conditions, torque was applied to push the wrist in the radial (positive, against gravity) direction (CF^+^: *τ*_*RUD*_ = 0.3 Nm) or in the ulnar (negative, with gravity) direction (CF^*−*^: *τ*_*RUD*_ =*−* 0.3 Nm). These CF conditions were included to provide a clearer interpretation of DF, as the effects observed during DF could result from either the independent control of individual muscles or a higher-order mechanism regulating co-contraction [6]. Specifically, CF controls for radial muscle activation (*u*_*R*_), primarily in response to CF^*−*^, and ulnar muscle activation (*u*_*U*_), primarily in response to CF^+^.

#### 1) Behavioral Study

EMG was used in the behavioral study only to assess muscle activity. We used Delsys Trigno Duo electrodes (sampling frequency: 2148 Hz), and the EMG signal was time-synced to task performance data (sampling frequency: 1000 Hz) via a common analog signal sent at the onset of each trial. EMG sensors were placed on the participant’s four major wrist muscles: flexor carpi radialis (FCR), flexor carpi ulnaris (FCU), extensor carpi ulnaris (ECU), and extensor carpi radialis (ECR). The forearm was prepared for electrode placement by shaving the arm hair and cleaning the skin with alcohol. Electrode placement was checked by observing large bursts of agonist muscle activity during flexion, extension, radial deviation, or ulnar deviation. A force-torque sensor (ATI mini40, ATI Industrial Automation, Apex, NC, USA) in the device handle was used to measure user-applied torques during the task.

#### 2) Neuroimaging Study

The neuroimaging study took place at the University of Delaware Center for Biomedical and Brain Imaging with a Siemens Prisma 3T scanner and 64-channel head coil. All functional images were acquired using a multiband 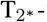 weighted EPI sequence (spatial resolution: 2 *×* 2 *×* 2 mm^3^, 60 slices, TE=30ms, TR=1s, acceleration factor=4, Image Size=[110 110 60]). We also acquired a T1-weighted anatomical image (1 *×* 1 *×* 1 mm^3^, TE=3.02ms, TR=2.3s, Image Size=[160 256 256]) for image registration and normalization and a GRE-field map (3 *×* 3 *×* 3 mm^3^, TE=4.92ms, TR=400ms) for unwarping due to magnetic field inhomogeneities. We monitored participants’ pulse and respiration with an MR-compatible pulse oximeter and respiration belt (Siemens Physiology Monitoring Unit).

### C. Protocol

After participants were positioned on the bed with the robot, they practiced the task in a transparent (NF) condition. During this time, they were instructed to practice making straight point-to-point movements within the cued time window. Participants were able to self-select when they felt comfortable enough with the task instructions and felt ready to end the practice session, after typically no more than 100 trials.

#### 1) Behavioral Study

After familiarization, participants performed six blocks of 200 trials each, with about five minutes of rest between blocks to reduce fatigue (Fig. 2). All but the final task block featured horizontal movements, and the final block featured vertical movements. The goal of the task was always the same, but the robot was commanded in different force modes across blocks. The first block was always an NF task to assess baseline task performance, followed by three randomly ordered DF, CF^+^, and CF^*−*^ blocks (Fig. 2). The final two blocks were added for EMG calibration, where participants were subject to viscous resistive forces during horizontal 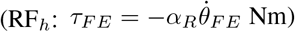 and vertical 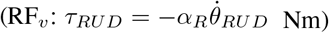 movements, where *α*_*R*_=0.050 Nm/deg. The RF tasks were intended to elicit strong agonist activation, improving the signal-to-noise ratio for improved EMG calibration.

**Fig 2.**
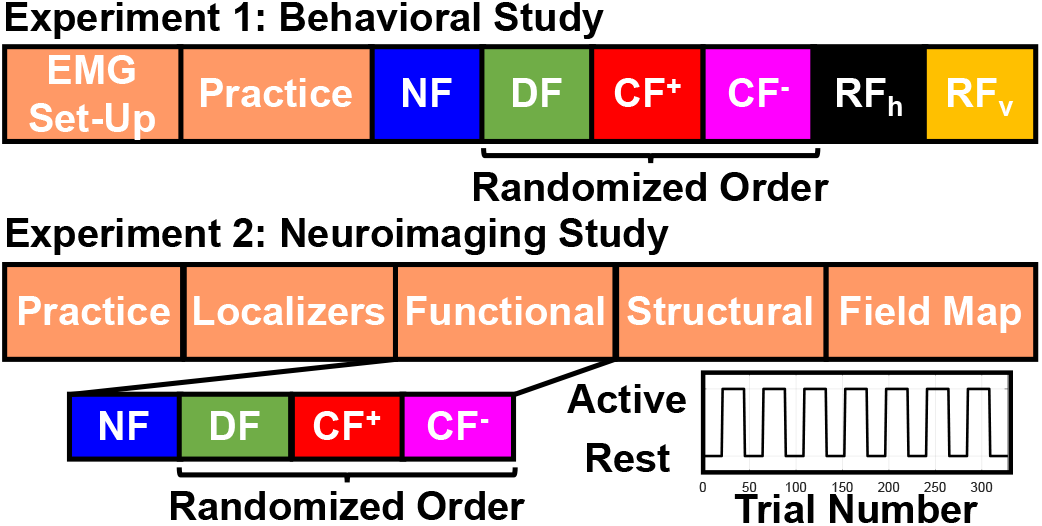
Experimental protocol of the behavioral and neuroimaging studies.

#### 2) Neuroimaging Study

In the neuroimaging study, participants performed four tasks (Fig. 2). Each task consisted of 168 active trials divided evenly into seven blocks, interspersed with 19.5 s passive blocks. During these passive blocks, the robot locked in its current position, and participants were instructed to watch the virtual cursor make straight point-to-point movements between the targets. This condition acts as our visual control to match the visual stimulus seen during task performance. Within each task, all active trials were controlled under the same force mode, which corresponded to the same conditions used in the first four blocks of the behavioral EMG study. The first block was always NF, followed by a random order of DF, CF^+^, and CF^*−*^.

### D. Data Analysis

#### 1) Trial Onset and Normalization

To consistently analyze data collected over multiple trials, we established three specific time windows: Early, Middle, and Late. The Early window spanned from 150 ms before trial onset (*t*_0_) and lasted until *t*_0_. In our data analysis, trial onset was defined as the instant when the cursor exceeded a position threshold of 1 deg away from the starting target in the direction towards the cued target. The Middle time window started at *t*_0_ and ended at the point of maximum velocity (*t*_*vmax*_). Finally, the Late time window covered the period from *t*_*vmax*_ until the end of the trial (*t*_*end*_), defined as the time when the cursor reached a lateral position within 1 deg of the ending target. Task kinematics, kinetics, and EMG were normalized via resampling so that the instants of *t*_*−*150_, *t*_0_, *t*_*vmax*_, and *t*_*end*_ coincided across participants.

#### 2) EMG-based Joint Torque and Stiffness Estimation

EMG signals were bandpass filtered at 30-500 Hz, rectified, and the envelope was taken via a 10 Hz low-pass, zero-shift 4th order Butterworth filter. Muscle moment arms (*ρ*) were obtained at each joint position using OpenSim [7].

For each muscle, we time-shifted the EMG signal (*m*) to synchronize with kinematic data, accounting for the electromechanical delay. We then related this delayed EMG signal (*m*(*t − t*_*d*_)) to muscle forces (*F*_*m*_) via a muscle-specific scaling factor (*γ*) (Eqn. 1). While this simplistic EMG-force relationship neglects the position and velocity-dependent effects of muscle activation, these effects may be negligible in our task because movements are small amplitude (range *<* 20 deg) and slow during the main region of interest (quasi-static in the Early window, *t*_*−*150_ : *t*_0_). Within this range, the peak active muscle force is predicted to change by 0.6-45%, as predicted by the force-length curves in the OpenSim wrist model [7]. We are primarily interested in relative differences across force conditions, where the muscle lengths and velocities are similar, so estimation errors across conditions should be comparable.

We calibrate *γ* at the joint level by relating task dynamics and predicted user-generated torques (*τ*_*pred*_) to the forcetorque sensor measured torques (*τ*_*meas*_) via linear regression (Eqn. 2-3). For calibration, we included all active trials across all six force conditions.

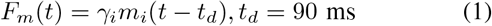

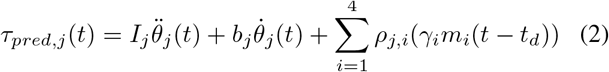

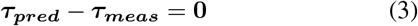

In these equations, *j* refers to the degree of freedom (FE or RUD) and *i* refers to the muscle (FCR, FCU, ECU, or ECR). Once *γ* was estimated for each muscle, we calculated human-generated torques in the radial direction only (*u*_*R*_), ulnar direction only (*u*_*U*_), and net RUD torque (*τ*_*RUD*_) as:

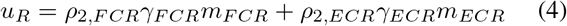

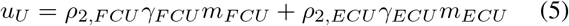

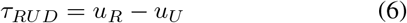

where *ρ*_2,*i*_ refers to the muscle moment arm in RUD.

Muscle stiffness is typically a function of its mechanical properties (i.e., stiffness of its cross-bridges), length, and cross-sectional area. However, it has been found experimentally that muscle stiffness within a small range can be directly related to force via a scaling constant *α/l*_0_, where *α*=23.4 [n.u.] and *l*_0_ is the optimal muscle fiber length at maximum activation [8]. We used this relationship to estimate muscle-level stiffness (Eqn. 7) and related it to joint-level stiffness via its moment arms (Eqn. 8), where ***J*** is the Jacobian matrix comprised of the muscle moment arms from OpenSim [7].

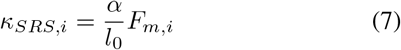

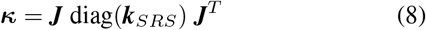

***κ*** reflects the force opposed by the participant when a unit displacement is applied, such that the magnitude of stiffness along a specific direction, *p*, is described as

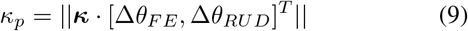

Thus, we extract RUD stiffness as *κ*_*RUD*_ = || ***κ***.[0, 1]^*T*^||.

Group-level differences were assessed using the average across all trials and analyzed using a linear mixed model. Significant force condition effects were followed by pairwise student’s t-tests, with significance set at *p <* 0.05. All statistics were performed using only the Early time window (*t*_*−*150_ : *t*_0_) to evaluate anticipatory motor commands, but the full trial is included for visualization.

#### 3) Statistical Analysis of Imaging Data

Neuroimaging data preprocessing was performed using SPM12 [9], following a standard pipeline: images were unwarped and field map corrected, then realigned, normalized into standard (MNI) space, and finally smoothed with an 8 mm Gaussian kernel.

The neural response was modeled as

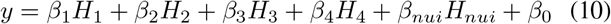

where H_1_, H_2_, H_3_, and H_4_ represent the task regressors for active blocks of NF, DF, CF^+^, and CF^*−*^ trials convolved with the standard hemodynamic response function. H_*nui*_ is a set of 25 nuisance regressors (18 physiological noise regressors, 6 head movement regressors, and 1 linear signal drift regressor). Physiological noise regressors were modeled using the RETROICOR algorithm via the TAPAS PhysIO Toolbox [10].

Analysis of the General Linear Model (GLM) (Eqn. 10) yielded maps of beta-weights, relating signal measured in each voxel to each regressor of interest (*β*_1_, *β*_2_, etc.). We report voxels that activate more in one condition relative to others, based on linear contrasts *c*_*i*_ of the extracted *β* coefficients. Due to the study’s limited number of participants, we report voxels with t-scores exceeding 4. This threshold was selected as a representative level for significance found in our prior work with larger study sizes [11], [12]. Activation regions were labeled using the Juelich Histological atlas for cortical regions [13], Harvard-Oxford atlas for subcortical regions [14], and MNI FNIRT atlas for the cerebellum [15].

For ease of notation, we define four activation maps to describe voxels related to the physiological phenomena of interest, i.e. areas of the brain involved in the control of force and impedance. Specifically, the K map identifies voxels that contribute to stiffness modulation, which we define as those whose BOLD signal is upregulated more during DF compared to NF (*c*_*K*_ = *β*_*DF*_ *− β*_*NF*_). The *u*_*R*_ and *u*_*U*_ maps refer to torque maps in the radial direction or ulnar direction, respectively. Voxels in the *u*_*R*_ and *u*_*U*_ maps respond more to the CF^*−*^ and CF^+^ compared to NF, such that 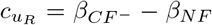 and 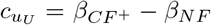. The *T* map isolates voxels specific to torque in any direction, such that 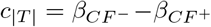, and both positive and negative statistical parametric coefficients are evaluated. Finally, the map CC relating to the neural control of co-contraction is extracted by identifying the set of voxels that were up-regulated in condition DF relative to baseline, compared to the sum of conditions CF^+^ and CF^*−*^, relative to baseline i.e., using the linear contrast (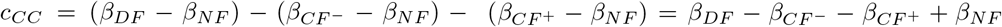)With this formulation, the intersection of the K and CC maps reflects voxels that tune to co-contraction.

## III. Results

### A. Behavioral Results

As confirmation of reasonable EMG to torque estimation, *τ*_*RUD*_ levels in the Early window (*t*_*−*150_ : *t*_0_) differed significantly across conditions in the expected directions (*p* = 0.002) (Fig. 3). The torque generated in response to CF^*−*^ was significantly greater than all other conditions (CF^*−*^ *−* NF: *p* = 0.002, CF^*−*^ *−* DF: *p* = 0.011, CF^*−*^ CF^+^: *p <* 0.001), and *τ*_*RUD*_ was not significantly different between NF and DF conditions (*p* = 0.263). The torque produced during CF^+^ was lower than both DF (*p* = 0.026) and CF^*−*^ (*p* = 0.002), but not significantly different than NF (*p* = 0.176).

**Fig 3.**
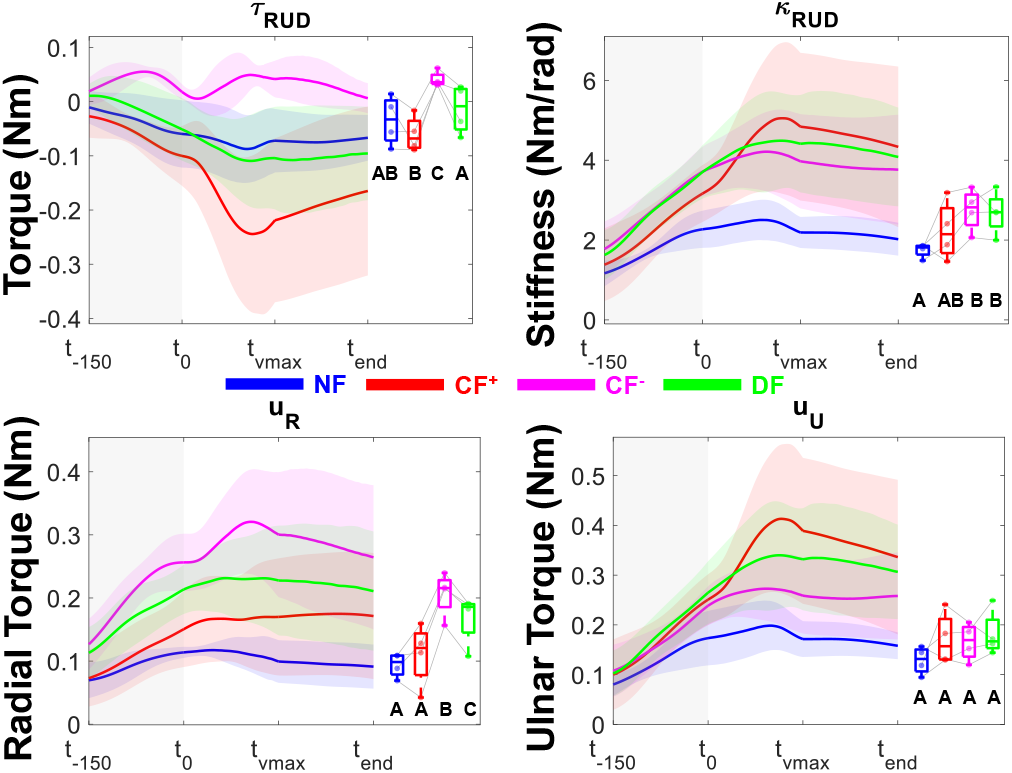
Group-level torque and stiffness estimates. The boxplots show the distribution of each subject’s mean activity in the shaded time window (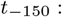 *t*_0_). Significance is expressed in the connecting letters format, where pairs of conditions that do not share a letter are significantly different.

The torque relationships in the radial direction (*u*_*R*_) follow a similar pattern (Fig. 3). The largest *u*_*R*_ magnitude occurred in the CF^*−*^ condition, significantly exceeding NF (*p <* 0.001), DF (*p* = 0.007), and CF^+^ (*p <* 0.001). There was no significant difference between NF and CF^+^ (*p* = 0.158). However, despite aiming to match *u*_*R*_ levels between DF and CF^*−*^ to test the muscle-level control hypothesis of co-contraction, *u*_*R*_ was significantly smaller in DF (*p* = 0.007).

For torque in the ulnar direction (*u*_*U*_), there were no significant differences across conditions (*p* = 0.218) (Fig. 3). The desired relationship would be to match *u*_*U*_ levels between DF and CF^+^ (where ulnar activation is agonist), and we are correctly not observing any significant difference between them. However, ideally we would like to see higher levels of *u*_*U*_ in CF^+^ compared to CF^*−*^ or NF to confirm that significant ulnar activation is necessary for the CF^+^ task. In its current state, the gravitational force may be canceling out much of the applied force, requiring only small ulnar activation.

Stiffness (*κ*_*RUD*_) was also significantly modulated by force condition (*p* = 0.020) (Fig. 3). While there was no significant difference in *κ*_*RUD*_ across pairs of the non-zero force conditions (CF^+^ *−* CF^*−*^: *p* = 0.097, CF^*−*^ *−* DF: *p* = 0.795, CF^+^ *−*DF: *p* = 0.148), *κ*_*RUD*_ was smaller in NF than CF^*−*^ (*p* = 0.006) and DF (*p* = 0.009). There was no significant difference between NF and CF^+^ (*p* = 0.115).

### B. Neuroimaging Results

Group-level fMRI results are shown in Fig. 4 and Table 1. Clusters of activation for stiffness modulation (K map) localized in the bilateral premotor cortices (Brodmann’s Area 6), cerebellum (left V and vermis VI lobules), and the thalamus and subthalamic region across both hemispheres, including the pallidum, amygdala, and caudate. We also observed some activation clusters in the occipital lobe. For the maps describing torque generation (*u*_*R*_, *u*_*U*_, and |*T*|), there were large regions of activation in the cerebellum, as well as the visual areas of the brain, such as the lateral occipital cortex and visual cortex V2 (Brodmann’s Area 18). The CC map, which describes the neural control of co-contraction, primarily resulted in active voxels in the bilateral premotor cortices, thalamus, and subthalamic regions (putamen, pallidum, and amygdala).

**TABLE 1.**
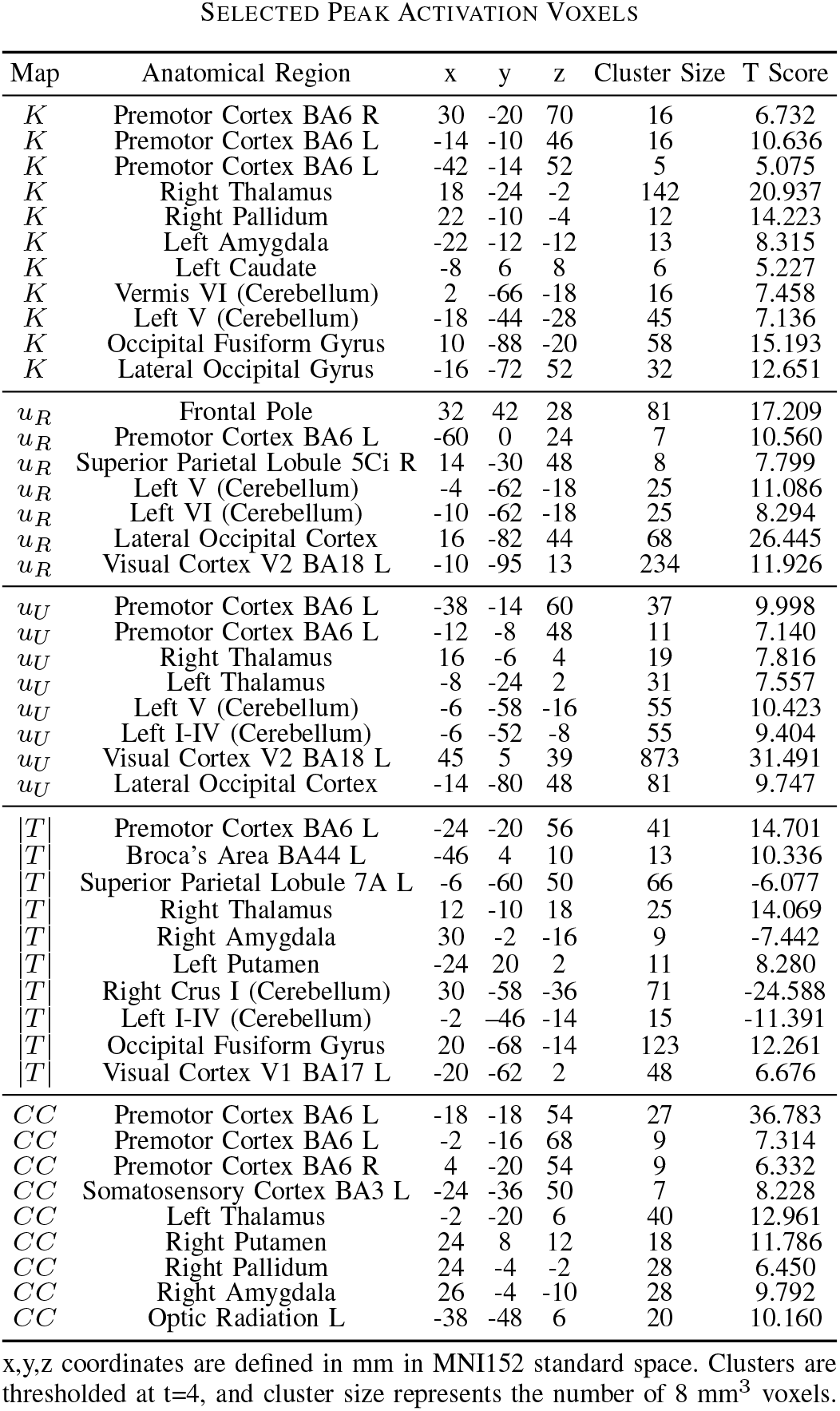
Selected Peak Activation Voxels.

**Fig 4.**
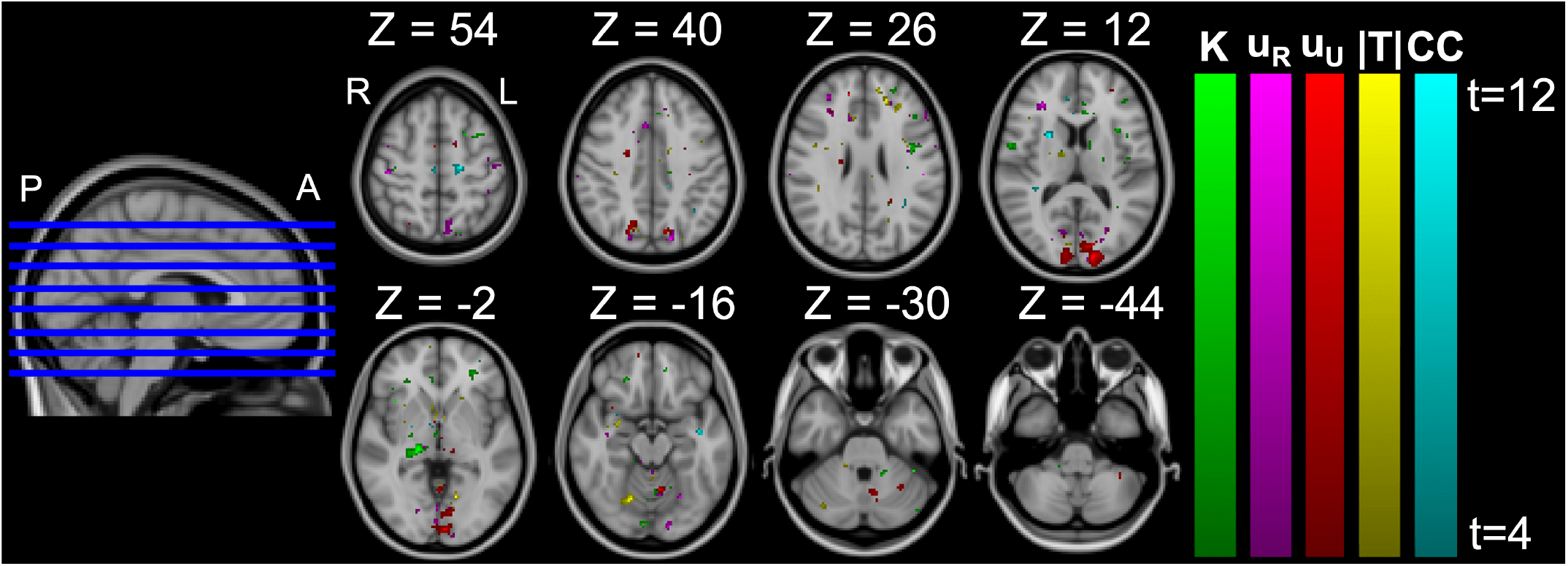
Group-level t-score activation maps for task-based regressors. These maps refer to contrasts *c*_*K*_ *>* 0, 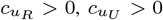, *c* ∣ *T* ∣ / = 0,and *c*_*CC*_ *>* 0. For visualization of the t-score gradients, each contrast was thresholded between t={4,12}. Z coordinates refer to axial slices in MNI152 coordinates in mm.

## IV. Discussion and Conclusion

The primary objective of this study was to examine the neural substrates associated with force and impedance control during dynamic wrist-pointing. We showed that our task significantly modulates stiffness (*κ*_*RUD*_) in DF relative to NF (Fig. 3), enabling the use of *c*_*K*_ to study stiffness modulation, as well as effects due to increased muscle activation. To isolate torque generation effects in the radial or ulnar direction, we included the constant force conditions in our paradigm to match agonist activation levels with those in DF. However, *u*_*R*_ was significantly greater in CF^*−*^ than DF, despite our aim to match levels. Additionally, CF^+^ produced a weak *u*_*U*_ response due to gravity reducing the applied force magnitude. This implies that our contrasts 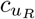 and 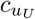 may not allow us to fully compensate for the effects of agonist activation. In a future study, we will increase the instability of DF to provide more stiffness modulation relative to the CF conditions and a better match of *u*_*R*_ between DF and CF^+^. We also plan to increase the magnitude of CF^+^ to overcome gravity. These changes should improve the relationships across conditions.

Our fMRI study revealed activation clusters within the cortico-thalamic-cerebellar network (Fig. 4, Table 1). Consistent with prior studies [16], clusters related to torque generation (*u*_*R*_, *u*_*U*_, and |*T*| maps) were observed in the dorsal premotor cortex. We also observed activation clusters in the ventral premotor cortex in the K, *u*_*R*_, *u*_*U*_, and |*T*| maps, supporting its role in voluntary co-contraction [16]. The overlap of the K and CC maps, representing regions specific to co-contraction control, identified only a few common voxels in the left thalamus, left amygdala, and anterior intra-parietal sulcus. We will need to recruit additional participants to understand if these areas are truly significant. Additionally, cerebellar activation in the K, *u*_*R*_, *u*_*U*_, and |*T*| maps aligns with the cerebellum’s proposed role in managing complex dynamics [17], [18].

We also observed large activation clusters in the visual cortex in the *u*_*R*_ and *u*_*U*_ maps. This activation may result from inadequate matching across conditions of the error feedback displayed at the end of each trial. Because error magnitude and direction differed across conditions (Fig. 5), the feedback cursor appeared at a different location on the screen, invoking a unique visual response. To address this for future studies, we plan to improve the visual control condition shown during rest by displaying errors for the virtual cursor that are representative of the actual errors of the participant during that condition. Additionally, we plan to incorporate explicit error regressors into the GLM (Eqn. 10).

**Fig 5.**
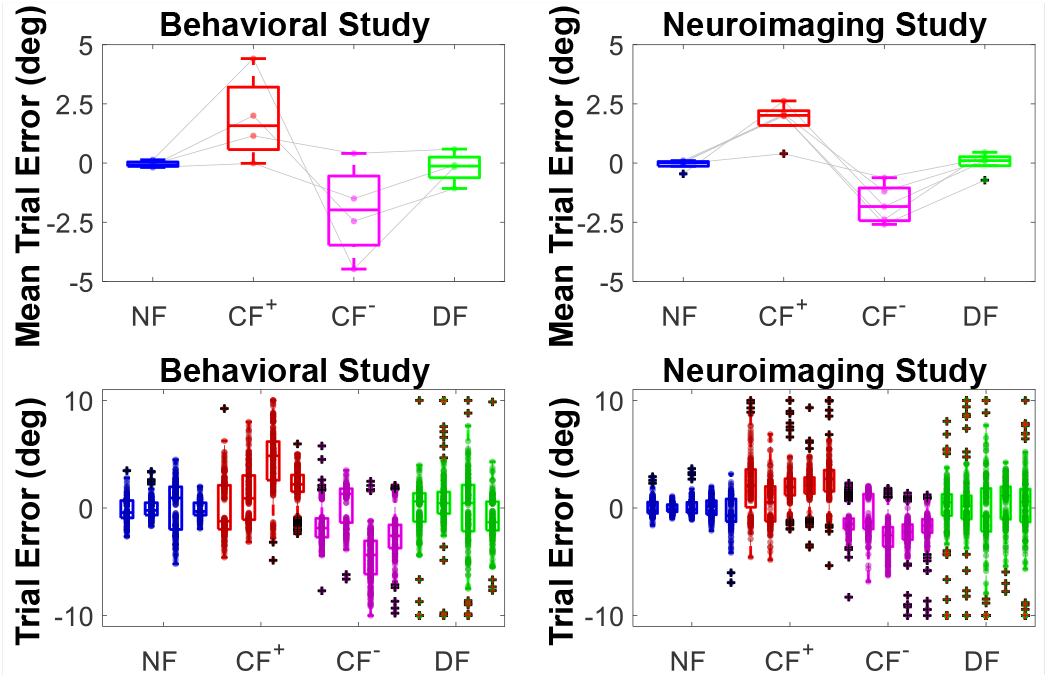
Trial errors displayed at the end of each trial. The top row shows the distribution of each subject’s average error in each force condition, and the bottom row shows the within-subject distribution of error across trials and conditions, where each boxplot represents one subject.

In conclusion, we demonstrated that the DF condition sufficiently modulates stiffness relative to NF, and our CF conditions properly modulate torque. However, while these relationships are sufficient to interpret joint-level neural activation, we need to refine the magnitudes of these conditions to fully control for muscle-specific activation (*u*_*R*_ and *u*_*U*_). Our fMRI study revealed activation within the cortico-thalamic-cerebellar network, with notable areas including the premotor cortex, thalamus, and cerebellum.

## Notes

This work is supported by the National Science Foundation under Award 194712 and the University of Delaware Graduate College through the Doctoral Fellowship for Excellence.

### Competing Interest Statement

The authors have declared no competing interest.

